# Stable *DNMT3L* Overexpression in SH-SY5Y Neurons Recreates a Facet of the Genome-Wide Down Syndrome DNA Methylation Signature

**DOI:** 10.1101/2020.11.09.374793

**Authors:** Benjamin I. Laufer, J. Antonio Gomez, Julia M. Jianu, Janine M. LaSalle

**Author notes:** The following authors contributed equally to the manuscript (Benjamin I. Laufer, J. Antonio Gomez).

## Abstract

Down syndrome (DS) is characterized by a genome-wide profile of differential DNA methylation that is skewed towards hypermethylation in most tissues, including brain. The molecular mechanisms involve the overexpression of genes related to DNA methylation on chromosome 21. Here, we stably overexpressed the chromosome 21 gene DNA methyltransferase 3L (*DNMT3L*) in the human SH-SY5Y neuroblastoma cell line and assayed DNA methylation at over 26 million CpGs by whole genome bisulfite sequencing at three different developmental phases (undifferentiated, differentiating, and differentiated). *DNMT3L* overexpression resulted in global CpG and CpG island hypermethylation as well as thousands of differentially methylated regions (DMRs). The *DNMT3L* DMRs were skewed towards hypermethylation and mapped to genes involved in neurodevelopment, cellular signaling, and gene regulation. Merging the DMRs into a consensus profile where the cell lines clustered by genotype and then phase demonstrated that different regions of common genes are affected. The hypermethylated DMRs from all pairwise comparisons were enriched for regions of bivalent chromatin marked by H3K4me3 as well as differentially methylated CpGs from previous DS studies of diverse tissues. In contrast, the hypomethylated DMRs from all pairwise comparisons displayed a tissue-specific profile enriched for regions of heterochromatin marked by H3K9me3 during embryonic development. Taken together, we propose a mechanism whereby regions of bivalent chromatin that lose H3K4me3 during development are targeted by excess DNMT3L and become hypermethylated, while excess DNMT3L also evicts DNMT3A from heterochromatin, resulting in hypomethylation. Overall, these findings demonstrate that *DNMT3L* overexpression during neurodevelopment recreates a facet of the DS DNA methylation signature.

## Introduction

Down syndrome (DS) is the leading genetic cause of intellectual disability and results from trisomy 21 (1). However, genes outside of chromosome 21 are also altered in DS and differences in gene expression and DNA methylation are observed across the entire genome. Most DS tissues exhibit differentially methylated sites that tend to be hypermethylated when compared to typically developing controls (2). Mechanistically, there are a number of genes located on chromosome 21 that belong to pathways related to DNA methylation and have the potential to result in the hypermethylation pattern observed in DS (2). Functional experimentation into the cause of hypermethylation has demonstrated a key role of the chromosome 21 encoded DNA methyltransferase *DNMT3L* at select genes (3). While DNMT3L is catalytically inactive, it is a regulatory factor that binds to and stimulates the *de novo* methyltransferases DNMT3A and DNMT3B (4–6).

Structurally, DNMT3L and DNMT3A form elongated heterotetramers through their C-terminal domains (DNMT3L-DNMT3A-DNMT3A-DNMT3L), and this complex multimerizes on DNA to form nucleoprotein filaments that spread DNA methylation over larger regions, such as CpG islands (7–9). Members of the *de novo* methyltransferase family (DNMT3A,B,L) contain an ATRX-DNMT3-DNMT3L (ADD) domain that binds to the unmodified histone H3 tail (H3K4me0), in order to localize DNA methylation to previously unmethylated regions, and this binding is inhibited by methylation at lysine 4 of H3 (H3K4me3)(10–12).

In mouse embryonic stem cells (mESCs) DNMT3L has been reported to be either a positive or negative regulator of DNA methylation depending on genomic context (13); however, another report found it to only function as a positive regulator (14). This difference appears to be due to the mESCs cell lines used as well as the methodologies used to assay DNA methylation, since the studies examined different regions of the genome at different levels of resolution. Experiments using transient *DNMT3L* overexpression in mESCs and somatic cells have demonstrated a role for DNMT3L in assembling a repressive chromatin modifying complex that silences retroviral sequences through *de novo* DNA methylation and methylation independent mechanisms (15). Finally, DNMT3L is also required for the establishment of parental imprinting patterns in developing gametes (16,17).

While *DNMT3L* is a prime candidate for the genome-wide DS DNA hypermethylation profiles, the genome-wide effect of *DNMT3L* overexpression has not yet been characterized. Previously, we profiled post-mortem DS brain and compared to matched controls using whole genome bisulfite sequencing (WGBS). We observed a genome-wide wide impact on DNA methylation profiles, where ∼75% of the differentially methylated regions (DMRs) were hypermethylated and mapped to genes involved in one-carbon metabolism, membrane transport, and neurotransmission. Given these observations, we sought to characterize the effect of DNMT3L overexpression on different phases of neuronal development by assaying differentiating SH-SY5Y cells.

SH-SY5Y cells are a human neuronal cell line that have been subcloned three times from the SK-N-SH cell line, which was derived from metastatic cells of a 4-year-old female with neuroblastoma. Unlike many other cancer lines, the SH-SY5Y cell line has a stable karyotype with a dominant ploidy of 2, which includes trisomy 1q and 7, gains on 2p and 17q, and losses on 14q and 22q (18–20). Similar to other cancer cell lines, SH-SY5Y cells have a relatively hypomethylated genome when compared to neurons from human post-mortem brain samples (21,22). SH-SY5Y cells display a mixed morphology that is primarily neuroblast-like with short processes; however, there is also a smaller proportion of epithelial-like cells, which likely reflects the cell lines multipotency and an origin from the neural crest. SH-SY5Y cells can be consistently differentiated from a neuroblast-like state into a homogenous population of mature dopaminergic neurons in 18 days (23). The differentiation protocol is a three phased reprogramming approach, which involves gradual serum deprivation and the addition of retinoic acid during the first two phases and then a transition to an extracellular matrix where the cells are given a neuronal media that lacks serum. The neuron-like cells become dependent on neurotrophic factors, such as BDNF, for their survival and the epithelial-like cells, which require serum, do not survive. Notably, during this differentiation process, serial splitting via a brief incubation with low strength trypsin allows for the selection of the less adherent neuron-like cells (for the next phase or harvest) and ultimately results in a morphologically pure culture of mature dopaminergic neurons in the final phase. Thus, SH-SY5Y cells serve as an ideal model system to investigate the methylome-wide effects of DNMT3L overexpression on undifferentiated, differentiating, and differentiated human neurons.

Here, we interrogated the contribution of DNMT3L, a chromosome 21 encoded gene, in DS-associated genome-wide changes in DNA methylation. Stable overexpression of DNMT3L in human SH-SY5Y cells provided a model system to test the effects of this known methylation regulator on genome-wide DNA methylation across neuronal differentiation. We found that DNMT3L overexpression recreated a facet of the genome-wide DNA profile observed in DS tissues, including brain. Interestingly, chromatin state analyses revealed that the DNMT3L-induced hypermethylated DMRs were enriched within bivalent domains, whereas the hypomethylated DMRs were enriched within heterochromatin. These results indicate that an increase in DNMT3L copy number during neuronal differentiation leads to chromatin-specific effects on DNA methylation.

## Methods

### Plasmids and Cloning

*DNMT3L* was cloned into an expression plasmid by Gibson assembly. First, human wild-type *DNMT3L* cDNA (IMAGE clone 1541874) was PCR amplified from pD3LMyc (4). PCR amplification primers were design with the complementary overhangs necessary for Gibson assembly using NEBuilder (New England Biolabs). The sequence for the forward primer was: TGTCTCATCATTTTGGCAAAATGGAGCAGAAGCTGATCTCAGAGGAGGAC. The sequence of the reverse primer was: TCACCGCATGTTAGCAGACTTCCTCTGCCCTCGCCACCTCCGCTGCCGCCTAAAGAGGAAGTGAGTTCTGTTGAAAAATACTTG. The reverse primer was designed to include a sequence coding for a self-cleaving P2A peptide inframe of DNMT3L. A PiggyBac-compatible expression plasmid, PbCAG-eGFP, was a gift from Joseph Loturco (Addgene plasmid # 40973) (24). PbCAG-eGFP was linearized with EcoRI. Both the linearized plasmid and the DNMT3L PCR amplicon were run on an agarose gel and purified in a silica column (Qiagen). Purified fragments were mixed at 1:3 molar ratio and assembled with NEBuilder HiFi DNA Assembly Mater Mix (New England Biolabs) according to the manufacturer’s instructions. Assembled products were then introduced into NEB Turbo E. coli competent bacteria by chemical transformation, plated on agar plates supplemented with ampicillin, and grown at 37°C according to manufacturer’s instructions. Colonies were then grown in LB and the plasmid was purified in spin silica columns according to manufacturer’s instructions. Resulting pb-DNMT3L-P2A-eGFP plasmids were confirmed by Sanger sequencing before proceeding to cell culture transfection experiments.

### Cell Culture and Transfection

Low passage SH-SY5Y cells in the undifferentiated growth phase were transfected with the plasmids via Lipofectamine 3000 (Invitrogen) according to the manufacturer’s instructions. After transfection, the cells were grown to a previously established timepoint when transient expression was shown to dissipate so that only cells with stable integration could be sorted for GFP via flow cytometry. After flow cytometry, the population of >95% GFP positive cells were re-cultured, expanded for ∼5 passages, and then differentiated according to Shipley *et al*. (23) except that the newer B-27 plus neuronal cell culture system (Gibco) was used and the retinoic acid was dissolved in DMSO. DNA was isolated from flash frozen cell pellets using the QIAamp DNA Micro kit (Qiagen) according to the manufacturer’s instructions.

### Whole Genome Bisulfite Sequencing (WGBS)

WGBS library preparation was performed using the post-bisulfite adaptor tagging (PBAT) method with the terminal deoxyribonucleotidyl transferase–assisted adenylate connector– mediated single-stranded-DNA ligation technique (25,26) via the Accel-NGS Methyl-Seq DNA Library Kit (Swift Biosciences) with the Methyl-Seq Combinatorial Dual Indexing Kit (Swift Biosciences) according to the manufacturer’s instructions. The library pool was sequenced across 2 lanes on an Illumina NovaSeq 6000 S4 flow cell for 150bp paired end reads to generate ∼150 million unique read-pairs (∼10X coverage) of the genome per sample.

### Bioinformatic Analyses

Trimming of adapters and methylation bias, screening for contaminating genomes, alignments to hg38, deduplication, calculation of coverage and insert size metrics, extraction of CpG methylation values, generation of genome-wide cytosine reports (CpG count matrix), and examination of quality control metrics were performed using CpG_Me (https://github.com/ben-laufer/CpG_Me) (27–30). DMR calling as well as most downstream analyses were performed using DMRichR, which utilizes the dmrseq and bsseq algorithms (https://github.com/ben-laufer/DMRichR) (30–32). Linear mixed effects model of the average methylation level were utilized for global and CpG island methylation level analyses, where genotype (3L or GFP) and cellular differentiation phase were direct effects, while cell line of origin was a random effect. Transcription factor motif enrichment testing using HOMER (33), DS pan-tissue enrichment testing via GAT (34), as well as enrichment testing for histone post-translational modifications (5 marks, 127 epigenomes) and chromatin states (15-state chromHMM model) (35,36) through LOLA (37), were all performed as previously described (30).

## Results

### Stable DNMT3L Overexpression in SH-SY5Y Cells

SH-SY5Y cells were transfected with either myc-tagged human DNMT3L (4) and enhanced GFP via a piggyBac transposase (24) construct or a matched control construct without DNMT3L (**Figure 1A**). GFP positive cells were selected by flow cytometry and then cultured again to enrich for a cell line with stable DNMT3L overexpression. For the experimental investigations, three developmental phases were assayed: 1) undifferentiated cells (growth), 2) the first phase of differentiation (phase 1), and 3) the final phase of differentiation (phase 3) that represents the pure and mature neuronal culture (**Figure 1B**). When compared to the growth phase, phase 1 neurons displayed a relatively elongated cell body with an increased number of longer neurites with neuronal growth cones that were starting to form synapses with their neighboring cells. Notably, phase 1 neurons are the most commonly assayed form of differentiated SH-SY5Y cells in the literature. Phase 2 is a relatively brief intermediate phase for the final selection that occurs in phase 3 and thus was not examined. After being plated as a monolayer, phase 3 neurons migrated to form large spheres of highly interconnected neurons, which were similar to neurospheres; however, they are were attached to an extracellular matrix and produced a 3-dimensional network with a larger scale organization that consists of long tracts of bundled processes. Notably, the GFP-only control cells displayed visibly higher fluorescence levels than those with DNMT3L + GFP, a result that was observed in all cell lines. Endogenous DNMT3L expression was confirmed by Western blot in all transfected cell lines and compared to a non-transfected cell line (WT). As expected, myc-tagged DNMT3L expression was only observed in the transgenic DNMT3L cell lines and not the WT or GFP-only control cell lines (**Figure 1C**).

**Figure 1:**
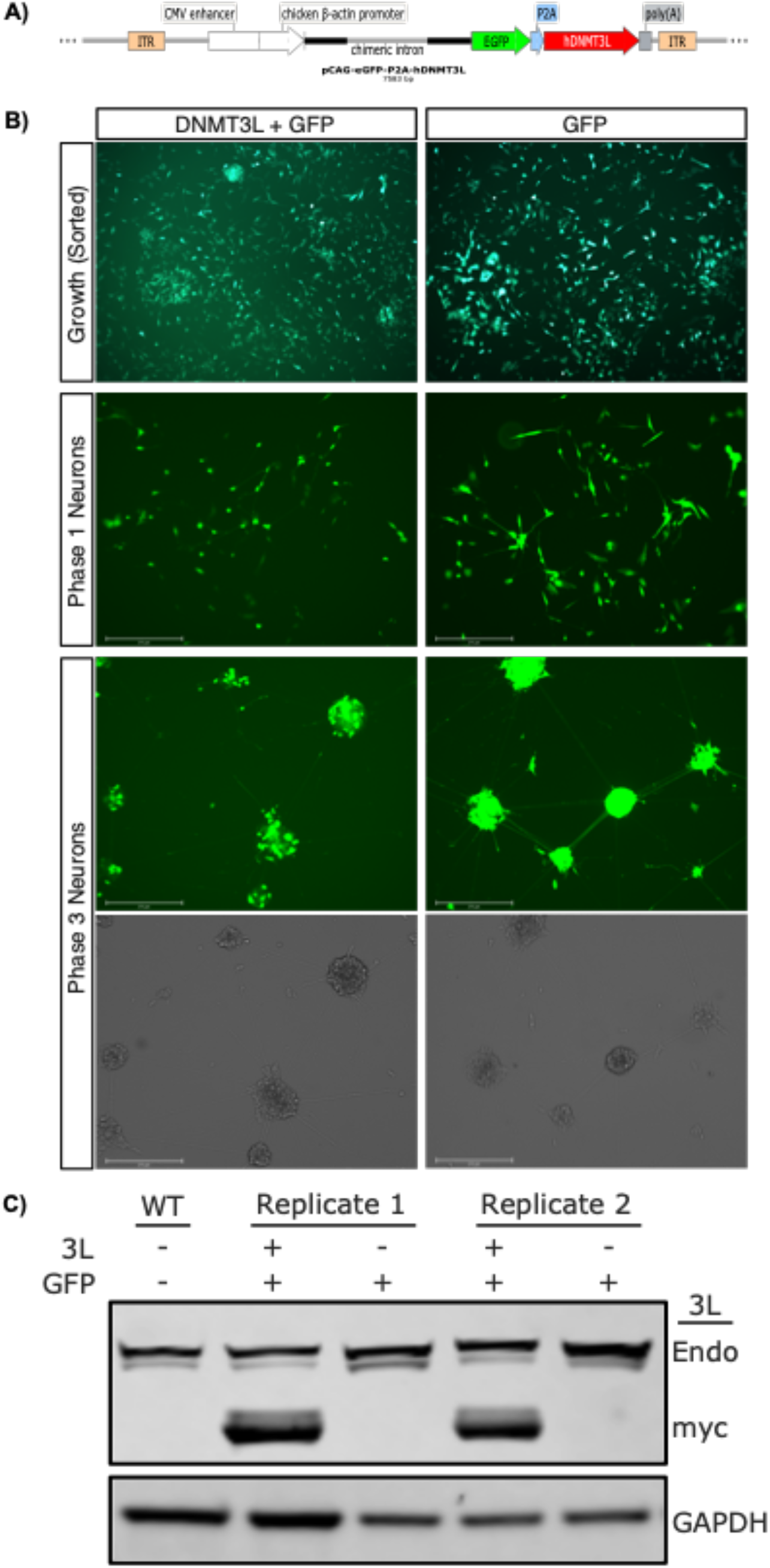
Stable overexpression of human *DNMT3L* in SH-SY5Y cells at different phases of neural differentiation. **A)** PiggyBac vector for stable integration of human *DNMT3L* (*hDNMT3L*), which was co-expressed with enhanced green fluorescent protein (eGFP). Upon translation the self-cleaving peptide (P2A) sequence between the polypeptide resulted in separate proteins. The vector was flanked by inverted terminal repeats (ITRs) that were recognized by the PiggyBac transposon. **B)** Representative live cell images the *DNMT3L* + eGFP and eGFP cell lines during the different phases assayed in either the GFP or phase comparison channels. **C)** Western blots of endogenous DNMT3L (Endo-3L), tagged transgenic DNMT3L (myc-3L), and GAPDH in wild-type SH-SY5Y cells (SH-WT), the DNMT3L + eGFP cell line (SH-3L), and the eGFP cell lines (SH-GFP).

### Stable DNMT3L Overexpression Results in Global and CpG Island Hypermethylation

Whole genome bisulfite sequencing (WGBS) libraries were constructed using DNA isolated from all cell lines and sequenced to >10x coverage, resulting in >26 million assayed CpGs. There was a significant (*p* = 0.016) increase in global CpG methylation (**Figure 2A**) and a significant (*p* = 0.004) increase in global CpG island methylation in cells with DNMT3L overexpression (3L) when compared to vector-only (GFP) controls. Cell developmental phase, herein referred to as “phase”, also had a significant (*p* = 0.004) effect on global CpG methylation and a significant (*p* = 0.030) effect on global CpG island methylation. In the growth phase, 3L cells showed 69.70% global methylation and 34.11% CpG island methylation, while the GFP cells showed 69.43% and 33.82%, respectively. In phase 1 of differentiation, 3L cells showed 70.38 % global methylation and 34.39% CpG island methylation, while GFP cells showed 69.93% and 33.89%, respectively. In phase 3 of differentiation, the 3L cells showed 69.68% global methylation and 33.95% CpG island methylation, while GFP cells showed 69.33% and 33.52%, respectively. Notably, both global CpG methylation and global CpG island methylation levels were the highest in phase 1 and the lowest in phase 3. Principal component analysis (PCA) revealed that the cells clustered together by phase for both the 20 Kb window and single CpG approaches (**Figure 2B**), but clustered by genotype for the CpG island approach (**Figure 2C**).

**Figure 2:**
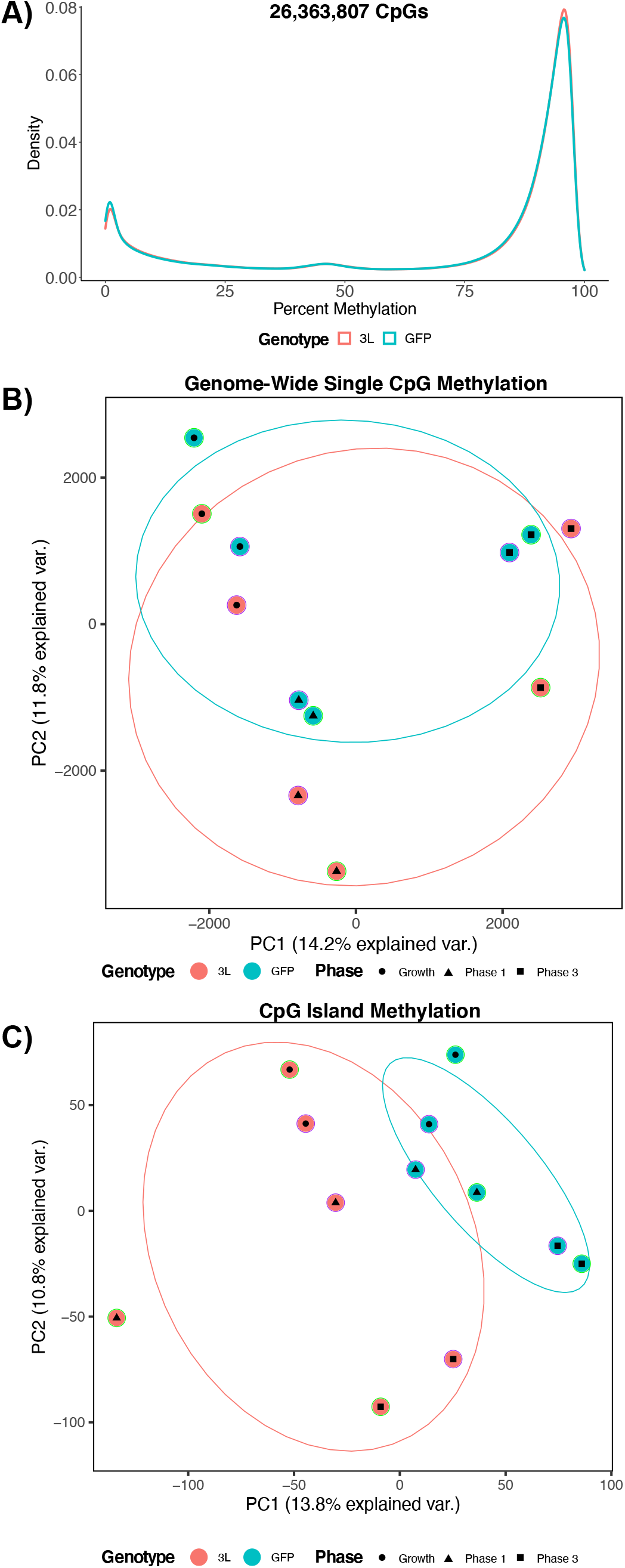
Global methylation profiles of *DNMT3L* overexpression. **A)** Density plots of the mean of smoothed individual CpG methylation values for each cell line at the different phases. **B)** Principal component analysis (PCA) of smoothed individual global CpG methylation values. **C)** PCA of smoothed individual CpG island methylation values. For each PCA, the color of the outermost shape represents the cell line, where green represents the first batch of cell lines and purple represents the second. The ellipses represent the 68% confidence interval, which represents 1 standard deviation from the mean for data with a normal distribution.

### DNMT3L DMRs Map to Genes Related to Neurodevelopment, Cellular Signaling, and Gene Regulation

In order to examine the specific genetic loci altered by DNMT3L overexpression during neuronal differentiation, differentially methylated regions (DMRs) were identified using pairwise genotype comparisons of all three phases. The replicate cell lines were utilized as biological replicates and batch was directly adjusted for in the analyses. The significant (*p* < 0.05) DMRs from each pairwise phase comparison were skewed towards hypermethylation (**Supplementary Table 1**). In the growth phase, 62% of the 6746 DMRs were hypermethylated, as well as 71% of the 5954 DMRs in phase 1, and 68% of the 6389 DMRs in phase 3 (**Figure 3**). DMRs were mapped to genes and gene ontology (GO) terms were slimmed to identify the least dispensable significant (*p* < 0.05) terms. Across all phases, when examining the effect of DNMT3L overexpression, these slimmed GO terms represented different neurodevelopmental processes, including synapse, signaling, and biological adhesion. However, there were also phase-specific GO terms, including the glucuronidation hierarchy of GO terms in the phase 1 comparison. While represented by xenobiotic glucuronidation, the glucuronidation hierarchy appears to represent retinoic acid, which is added to the media in phase 1, since the glucuronidation of retinoic acid is involved in neurodevelopment (38).

**Figure 3:**
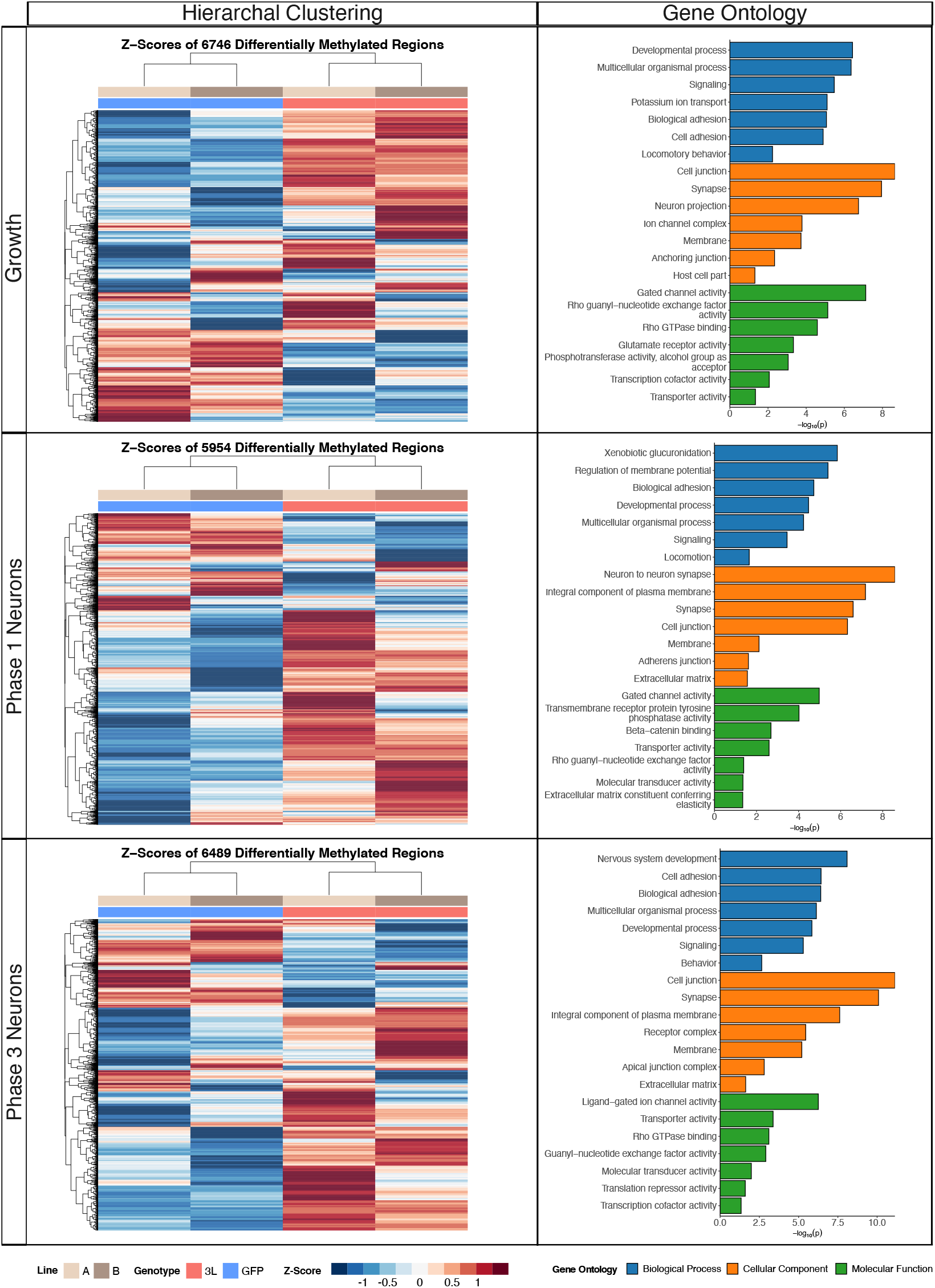
Differentially methylated region (DMR) profiles and slimmed gene ontology (GO) enrichments for *DNMT3L* overexpression at the different phases of neural differentiation. The heatmaps represent hierarchal clustering of Z-scores, which represent the number of standard deviations from the mean of the non-adjusted individual smoothed methylation value for each DMR. The bar plots represent the least dispensable significant enrichments (*p* < 0.05) slimmed GO.

To further test the functional relevance of DNA methylation changes resulting from DNMT3L overexpression, we tested the DMRs for enrichments within genomic regions annotated as CpG islands, CpG shores, CpG shelves, or open sea, as well as for enrichments within promoter and gene body annotations. Both the hypermethylated and hypomethylated DMRs from all pairwise phase comparisons were significantly (*q* < 0.05) enriched within CpG islands (**Figure 4A**) and significantly de-enriched within intergenic regions (**Figure 4B**). Furthermore, the hypermethylated DMRs from all pairwise phase comparisons were significantly (*q* < 0.05) enriched within promoters and regions of the gene body, while the hypomethylated DMRs from all pairwise phase comparisons were significantly (*q* < 0.05) enriched within introns.

**Figure 4:**
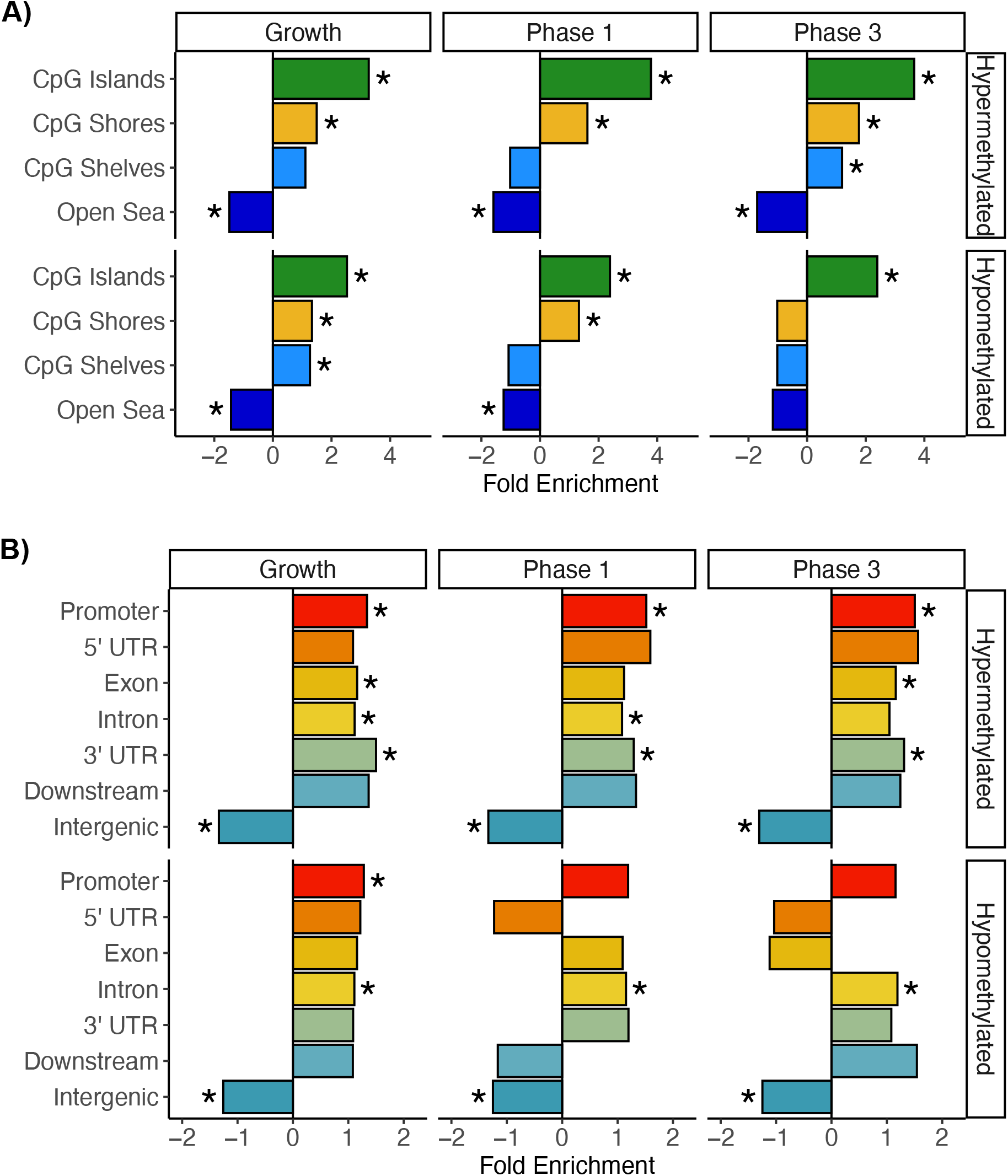
Annotation enrichments for DMRs from the growth, phase 1, and phase 3 comparisons. **A)** CpG and **B)** genic annotation enrichments for hypermethylated and hypomethylated DMRs. * = *q* < 0.05.

In order to understand the similarities and differences between the impact of DNMT3L overexpression on each developmental phase, DMRs from all pairwise comparisons were compared for overlap by both genomic coordinate and gene association. Overlaps of the DMRs by gene symbol included 962 in common to all three phases (**Figure 5A**), even though none overlapped by genomic coordinate in all three phases. The DMRs from all phase comparisons were then merged by sequence overlap to produce a consensus DMR profile, and hierarchal clustering analysis revealed that the cells clustered primarily by genotype and then developmental phase, and not by cell line of origin (**Figure 5B**). The consensus DMR profile highlighted not only the overall skew towards DMRs hypermethylation, but also clusters of DMRs that were specific to the differentiation phase. A meta p-value of analysis of the GO terms from all pairwise phase comparisons, revealed that the least dispensable significant (*p* <0.05) terms were largely similar to the individual comparisons and primarily represented neurodevelopment (**Figure 5C**).

**Figure 5:**
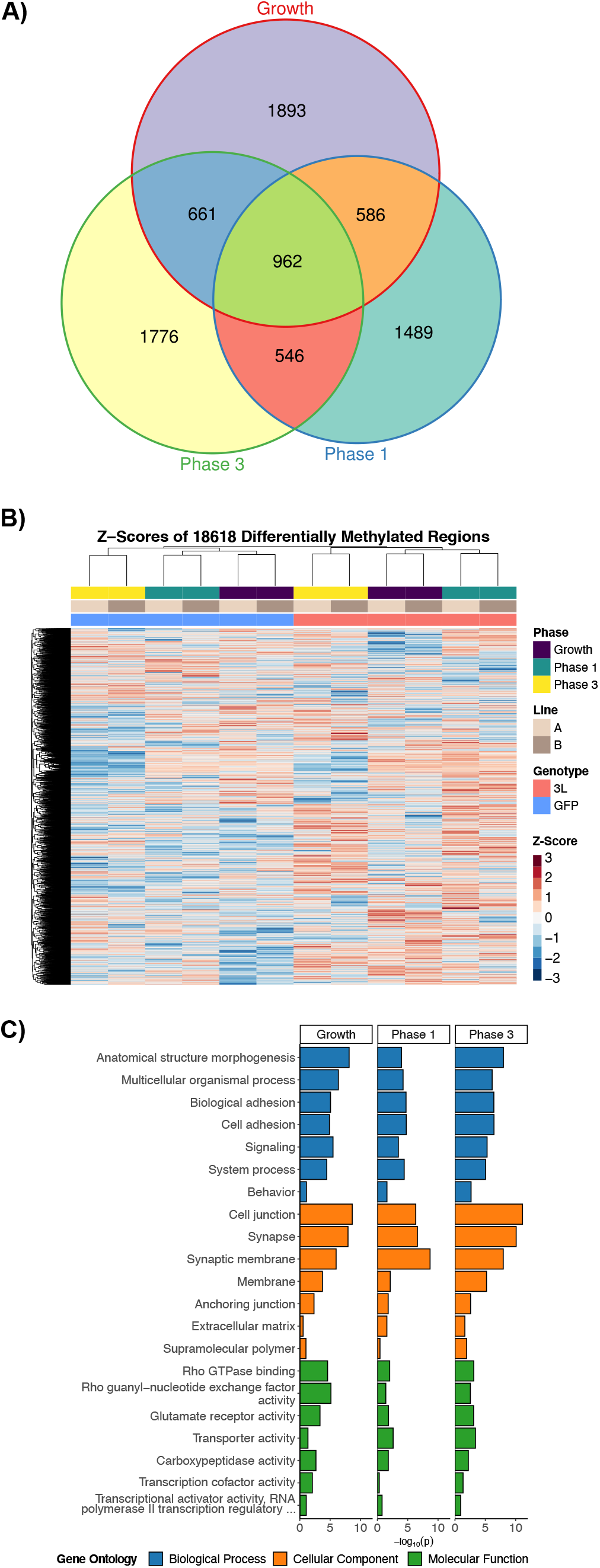
Consensus DMR profiles for *DNMT3L* overexpression across all phases of neural differentiation. **A)** Euler diagram of gene symbol overlaps for the genotype comparison at each phase of neural differentiation. **B)** Heatmap of hierarchal clustering of Z-scores for the consensus DMRs that are derived from merging the DMRs from each phase comparison by sequence overlap. **C)** Bar plot of comparison specific p-values from the meta-analysis of the least dispensable significant slimmed GO enrichments (*p*_*meta*_< 0.05).

### The Hypermethylated DMRs are Enriched for Bivalent Chromatin and DS Pan-tissue Regions

In order to further investigate the functional relevance between the hypermethylated and hypomethylated DNMT3L DMRs, enrichment analyses within chromatin state maps from the reference epigenomes were performed. The most significant (*q* < 0.05) chromHMM chromatin state enrichments overall were for the hypermethylated DMRs in regions of bivalent chromatin (**Figure 6A**) marked by H3K4me3 (**Figure 6B**) in stem cells. This effect was most pronounced in the differentiated cells. In contrast, the hypomethylated DMRs were most significantly (*q* < 0.05) enriched within heterochromatic regions (**Figure 6A**) marked by H3K9me3 (**Figure 6B**) in ESCs.

**Figure 6:**
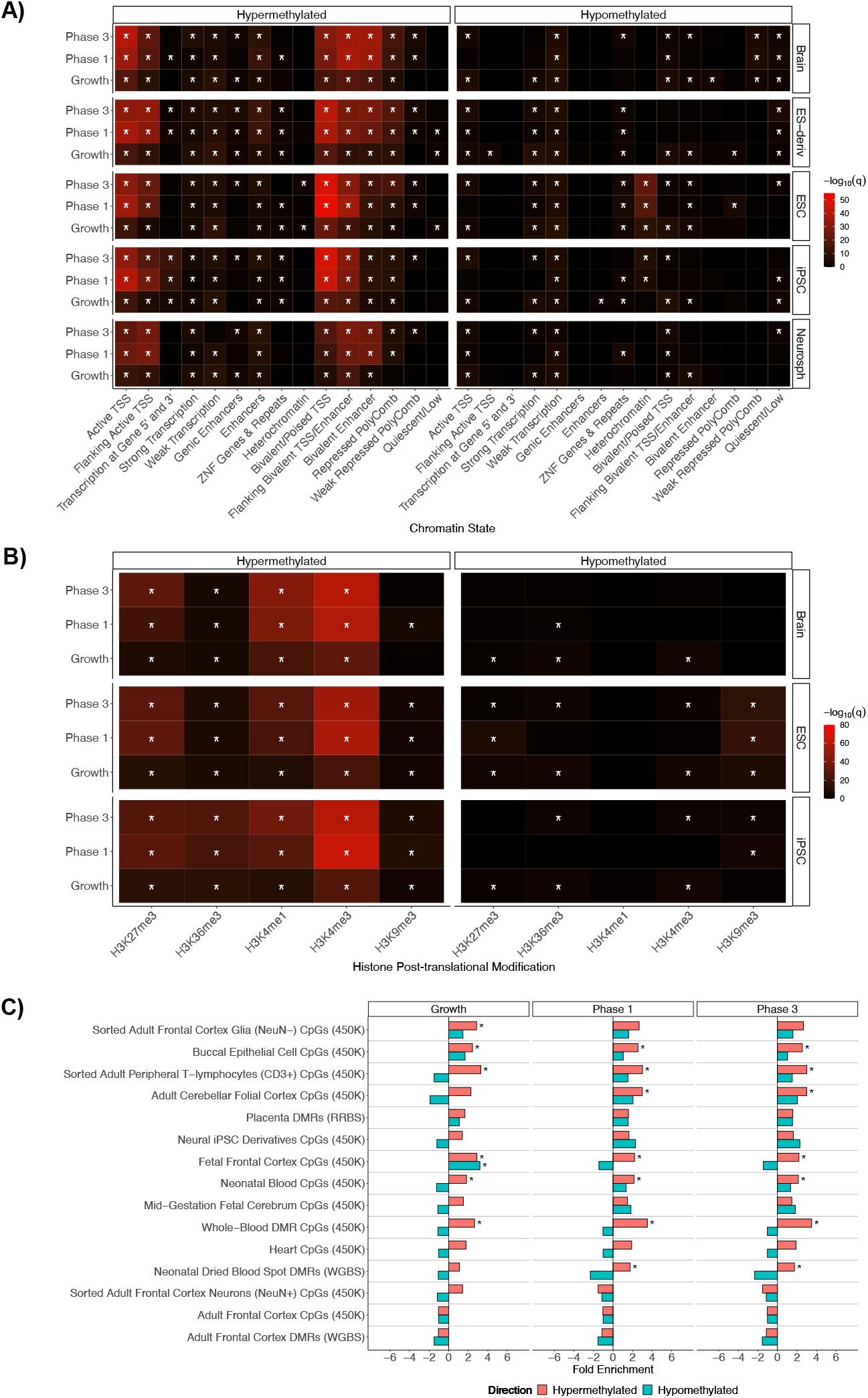
Reference epigenome and cross-tissue Down syndrome enrichment analyses for the hypermethylated and hypomethylated DMRs from the genotype comparisons at each phase of neural differentiation. **A)** Summary heatmap of top *q*-values for the chromHMM core 15-state enrichment analyses for the brain, embryonic stem cell derivatives (ES-derivatives), embryonic stem cells (ESC), induced pluripotent stem cells (iPSC), and neurospheres (Neurosph) categories. **B)** Summary heatmap of *q*-values for Roadmap epigenomics 127 reference epigenomes 5 core histone modification enrichment analyses for the Brain, ESC, and iPSC categories. **C)** Bar plot of Down syndrome cross-tissue analysis. All enrichments are relative to background regions. * = *q* < 0.05.

To determine the similarities between the DNMT3L-specific DMRs and the DNA methylation changes previously observed in DS, a pan-tissue comparison with differentially methylated DS sites across diverse tissues was performed (30,39–47). These results revealed significant overlap with DS differentially methylated sites, predominantly within the hypermethylated DNMT3L DMRs (**Figure 6C**). The hypermethylated DMRs from all pairwise phase comparisons were significantly (*q* < 0.05) enriched across a variety of tissues. Notably, the only significant (*q* < 0.05) enrichment for the hypomethylated DMRs was from the growth comparison and within fetal frontal cortex CpGs.

Also present were subsets of the 25 previously known pan-tissue and multi-tissue DS genes (2), which are all hypermethylated. This observation is in line with the previous observation that 24 out of the 25 pan-tissue and multi-tissue DS genes are hypermethylated. The growth comparison contained DMRs mapping to *ZNF837, RYR1, BCL9L, VPS37B*, and *RUNX1*. The phase 1 comparison contained a DMR mapping to *TEX14*. The phase 3 comparison contained DMRs mapping to *BCL9L, RFPL2, RYR1, ADAMTS10, MZF1*, and *RUNX1*.

While the DNMT3L DMRs overlapped with many of the 106 imprinted genes in the human genome, these results were not significantly higher than expected in enrichment analyses. The growth DMRs mapped to 18 imprinted genes: *SNRPN, ZNF597, ZFAT, NTM, LIN28B, GRB10, MIMT1, KCNQ1, ADTRP, ANO1, OSBPL5, FAM50B, ATP10A, SNORD116, MIR296, GLIS3, PPP1R9A*, and *KCNK9*. The phase 1 DMRs mapped to 19 imprinted genes: *ATP10A, MKRN3, PHLDA2, RB1, MESTIT1, IGF2, DGCR6L, ZFAT, NTM, MIR296, MAGI2, ZIM2, SNORD116, KCNK9, ZDBF2, NAA60, KCNQ1, GNAS*, and *KCNQ1OT1*. The phase 3 DMRs mapped to 22 imprinted genes: *OSBPL5, TP73, MKRN3, WT1, KCNK9, NAA60, PPP1R9A, ATP10A, SNRPN, PLAGL1, GRB10, SNORD116, SLC22A3, MEG8, MEG3, ZIM2, MIMT1, GNAS, UBE3A, NDN, IGF2*, and *ADTRP*.

### External Replicate Cell Line

To test the reproducibility of the observed effects on DNMT3L expression on genome-wide DNA methylation, we carried out a replication analysis using a separate lot of SH-5YSY cells. Like the two primary replicate cultures in our study, the third replicate was also derived from low passage cells from ATCC. DNMT3L overexpression was confirmed in the third replicate (**Supplementary Figure 1**). The main analyses were then repeated to include the third replicate. Similar to our original observation, there was still a significant (*p* = 0.002) increase in global CpG methylation (**Supplementary Figure 2A**) and a significant (*p* = 0.0008) increase in CpG island methylation from 3L overexpression. Phase had a significant (*p* = 0.01) effect on global CpG methylation but, in contrast to the previous analyses, there was no significant (*p* = 0.25) effect of phase on global CpG island methylation. Similar to the other analyses, PCA revealed that the cells clustered together by phase for both the 20 Kb window and single CpG approaches (**Supplementary Figure 2B**), but clustered by genotype for the CpG island approach (**Supplementary Figure 2C**). When the third replicate was included in the DMR calling (**Supplementary Table 2**), there was a stronger hypermethylation skew: 75% of the 3359 growth DMRs, 86% of the 4885 phase 1 DMRs, and 91% of the 7279 phase 3 DMRs (**Supplementary Figure 3**). The GO terms were also similar and the DMRs mapped to genes involved neurodevelopment, cellular signaling, and gene regulation (**Supplementary Figure 3**). The significant (*q* < 0.05) enrichment of the hypermethylated DMRs within CpG islands was also replicated; however, the hypomethylated DMRs were not significantly enriched for within CpG islands (**Supplementary Figure 4A**). Similar to the previous comparisons of just the two replicates, the hypermethylated DMRs from all pairwise comparisons were significantly (*q* < 0.05) enriched within promoters and regions of the gene body, while the hypomethylated DMRs from all pairwise comparisons were significantly (*q* < 0.05) enriched within introns (**Supplementary Figure 4B**).

From all of the pairwise phase comparisons, 711 of the DMRs overlapped by gene symbol (**Supplementary Figure 5A**). In the consensus DMR heatmap, the 3^rd^ replicate clustered closer with itself than by genotype, suggesting that the primary differences are due to the characteristics of the cell line before transfection (**Supplementary Figure 5B**). However, a heatmap of the consensus DMRs with just the 3^rd^ replicate showed the expected clustering by differentiation timepoint (**Supplementary Figure 5B**). The meta p-value GO analysis also showed similar terms related to neurodevelopment, cellular signaling, and gene regulation (**Supplementary Figure 5C**).

The chromHMM chromatin state and histone modification enrichments also replicated, although with a larger effect size. The hypermethylated DMRs were significantly (*q* < 0.05) enriched in regions of bivalent chromatin (**Supplementary Figure 6A**) marked by H3K4me3 (**Supplementary Figure 6B**) in stem cells, an effect most pronounced in the differentiated cells. The hypomethylated DMRs were most significantly (*q* < 0.05) enriched for within regions heterochromatin (**Supplementary Figure 6A**) marked by H3K9me3 (**Supplementary Figure 6B**) in ESCs. While the DS pan-tissue analysis also showed more significant (*q* < 0.05) enrichments for the hypermethylated DMRs in the replication analysis, the hypomethylated fetal cerebral cortex enrichment was no longer present in the growth comparison (**Supplementary Figure 6C**). Overall, the 3^rd^ replicate clustered differently from the other two cell lines and had a more pronounced hypermethylation profile; however, it still replicated the main conclusions of the primary analyses.

## Discussion

The results of this study confirmed previous observations and provided novel findings that are relevant to understanding the role of DNMT3L in establishing DNA methylation profiles during development and in DS. DNMT3L overexpression in undifferentiated, differentiating, and differentiated SH-SY5Y neurons resulted in a global increase in CpG and CpG island methylation. The DMRs mapped to genes involved in neurodevelopment, cellular signaling and gene regulation. Both DNMT3L genotype and cell differentiation phase were inter-related in the consensus DMR clustering, demonstrating that different regions of the same genes are affected during development. However, our results also highlight a directional dichotomy in the DNA methylation changes associated with DNMT3L overexpression. The hypermethylated DMRs showed a pan-tissue DS profile, in contrast to the hypomethylated DMRs that showed a distinct profile related to early neurodevelopment. There were also distinct differences in the chromatin state enrichments for the hypermethylated and hypomethylated DMRs.

The hypermethylated DMRs were enriched within regions of bivalent chromatin marked by H3K4me3. Bivalent chromatin consists of regions marked with both activating H3K4me3 and repressive H3K27me3 and serves to regulate key developmental genes in embryonic stem cells by repressing them during pluripotency and poising them for rapid activation upon removal of H3K27me3 during differentiation (48,49). Notably, while bivalent promoters in embryonic stem cells are unmethylated or hypomethylated, upon differentiation some bivalent promoters lose H3K4me3 and gain DNA methylation (14,50,51). Since DNMT3L recognizes unmethylated H3K4 (10), a possible mechanism is that regions of bivalent chromatin lose H3K4me3 as they differentiate and become hypermethylated by the excess DNMT3L that recognizes the H3K4me0. This hypothesis is consistent with the observation that phases 1 and 3 DNMT3L DMRs show much stronger enrichments within regions marked by H3K4me3 when compared to the growth phase. This proposed mechanism is also consistent with the growth phase still showing enrichment within regions marked by H3K4me3, as the cells are already committed to the neural crest lineage. Additionally, while bivalent chromatin is generally hypomethylated in normal cells, in cancer cells bivalent chromatin becomes hypermethylated and is associated with increased developmental gene expression (52,53). In neural progenitors, many promoters with the H3K27me3 modification gain DNA methylation during differentiation (54).

In comparison to the hypermethylated DMRs, the hypomethylated DNMT3L DMRs in our study were enriched within regions of heterochromatin marked by H3K9me3. Notably, the observation of DNMT3L causing hypomethylation has also been previously established in multiple studies. Knockdown of DNMT3L in mESCs resulted in 14,107 regions showing a decrease in methylation and 5,724 showing a gain in methylation (13). This ratio is comparable to our observations and the regions identified significantly (*p* = 0.0002) overlap with the genomic coordinates for the consensus DNMT3L DMRs. Mechanistically, DNMT3L is known to either release or sequester DNMT3A from heterochromatin, which enables preferential targeting of euchromatin (55). Taken together, we hypothesize that in our study, the release or sequestering of DNMT3A from heterochromatin by excess DNMT3L resulted in the preferential targeting of bivalent chromatin within regions that recently lost H3K4me3 as the cells differentiated.

While *DNMT3L* overexpression during neuronal differentiation recapitulates an important facet of the epigenetic signature of DS, it is not the only mechanism. The other methylation differences observed in DS brain are likely related to an increase in copy number of the other proteins related to DNA methylation and one-carbon metabolism that are also located on chromosome 21 as well the contributions of age and medication (2). The results also highlight that while the impacts of the methylation profile are reproducible, there is variation within the replicate cell lines that may be due to differences in integration sites, copy number, and lot of cells. Overall, our findings suggest that DNMT3L overexpression in neurons recrates a facet of the Down syndrome DNA methylation profile by targeting specific histone post-translational modifications and chromatin states that are related to neurodevelopment.

## Acknowledgements

This work was supported by a National Institutes of Health (NIH) grant [3R01ES015359-10S1] to JML, a Canadian Institutes of Health Research (CIHR) postdoctoral fellowship [MFE-146824] to BIL, a CIHR Banting postdoctoral fellowship [BPF-162684] to BIL, a Jane Coffin Childs Memorial Fund for Medical Research fellowship to JAG, a NIH T32 Fellowship to JAG [5T32CA108459], and the UC Davis Intellectual and Developmental Disabilities Research Center (IDDRC) [P50HD103526]. The flow cytometry was carried out by the UC Davis Flow Cytometry Shared Resource Laboratory and was supported by the UC Davis Comprehensive Cancer Center Support Grant (CCSG) awarded by the National Cancer Institute [NCI P30CA093373]. The library preparation and sequencing was carried out by the DNA Technologies and Expression Analysis Cores at the UC Davis Genome Center and was supported by a NIH Shared Instrumentation Grant [1S10OD010786-01].

## Authorship Contributions

BIL, JAG, and JML designed the study. JML acquired funding and supervised the project. JAG designed, cloned, and validated the constructs. JAG and BIL performed the transfections. BIL performed the cell culture differentiation and microscopy. JAG and JMJ performed the western blots. BIL performed the bioinformatic analyses. BIL interpreted the results and wrote the manuscript with intellectual contributions and edits from JAG and JML. All authors reviewed and approved the final manuscript.

---

**Supplementary Figure 1:**
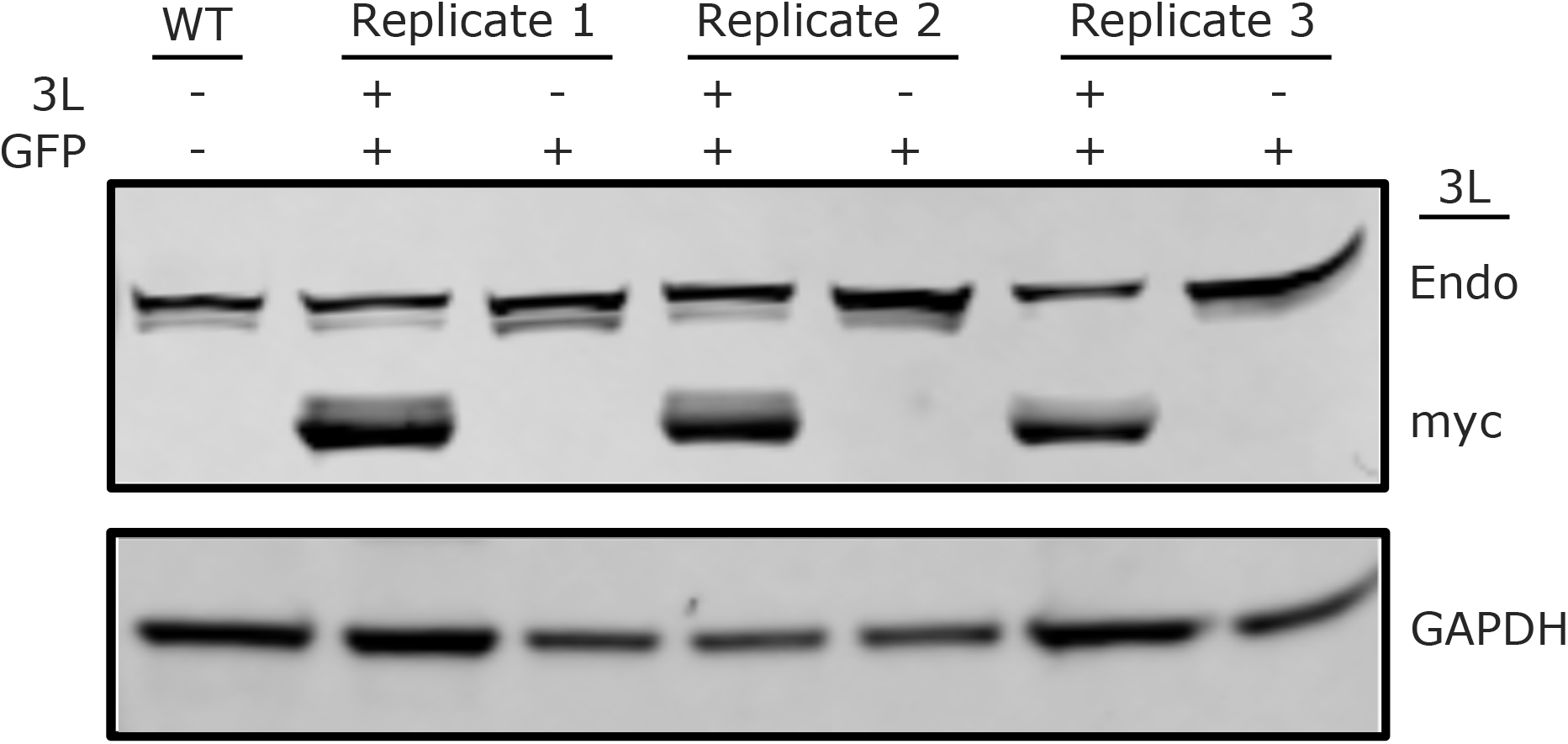
Western blots of endogenous DNMT3L (endo-3L), the tagged transgenic DNMT3L (tg-3L), and GAPDH in wild-type SH-SY5Y cells (SH-WT), the DNMT3L + eGFP cell line (SH-3L), and the eGFP cell line (SH-GFP) for all 3 cell line replicates.

**Supplementary Figure 2:**
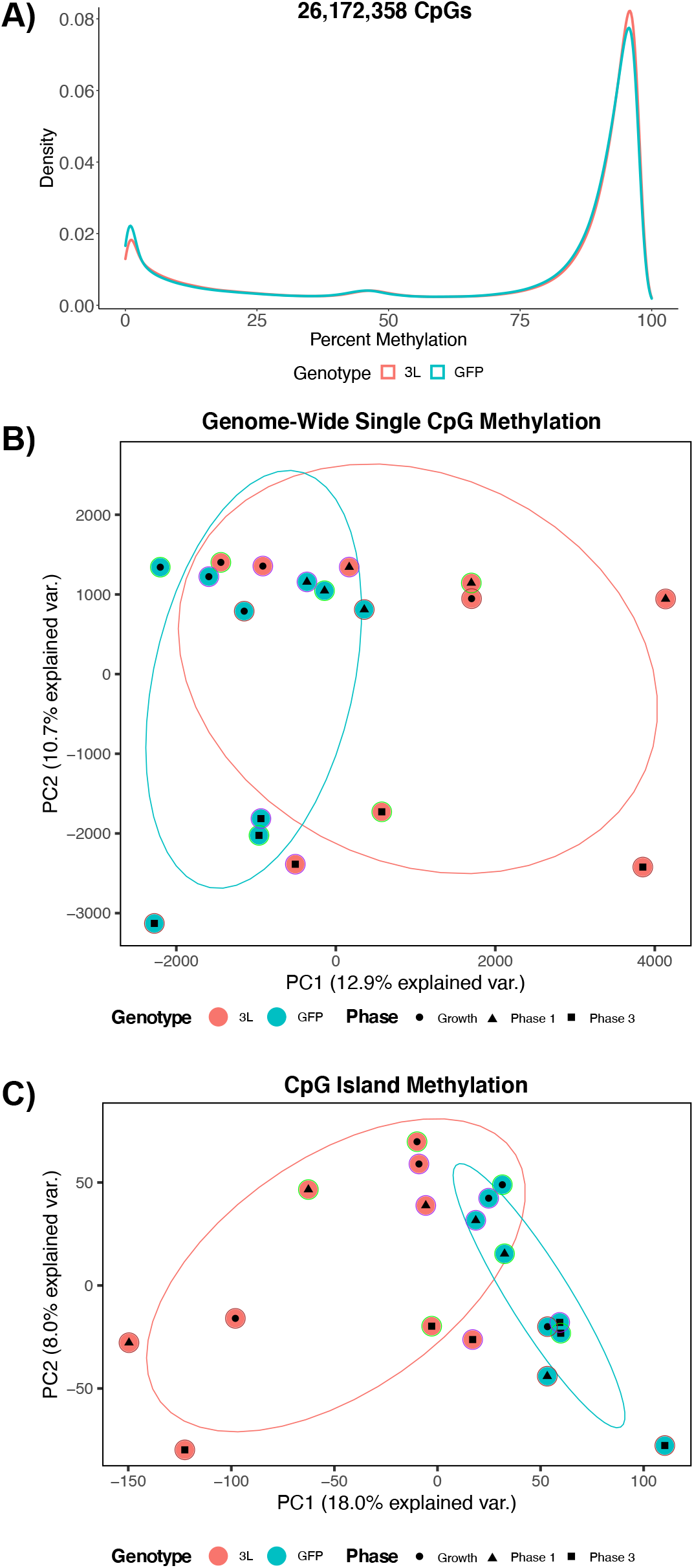
Global methylation profiles of *DNMT3L* overexpression for all 3 cell line replicates. **A)** Density plots of the mean of smoothed individual CpG methylation values for each cell line at the different phases. **B)** Principal component analysis (PCA) of smoothed individual global CpG methylation values. **C)** PCA of smoothed individual CpG island methylation values. For each PCA, the color of the outermost shape represents the cell line, where green represents the first batch of cell lines and purple represents the second. The ellipses represent the 68% confidence interval, which represents 1 standard deviation from the mean for data with a normal distribution.

**Supplementary Figure 3:**
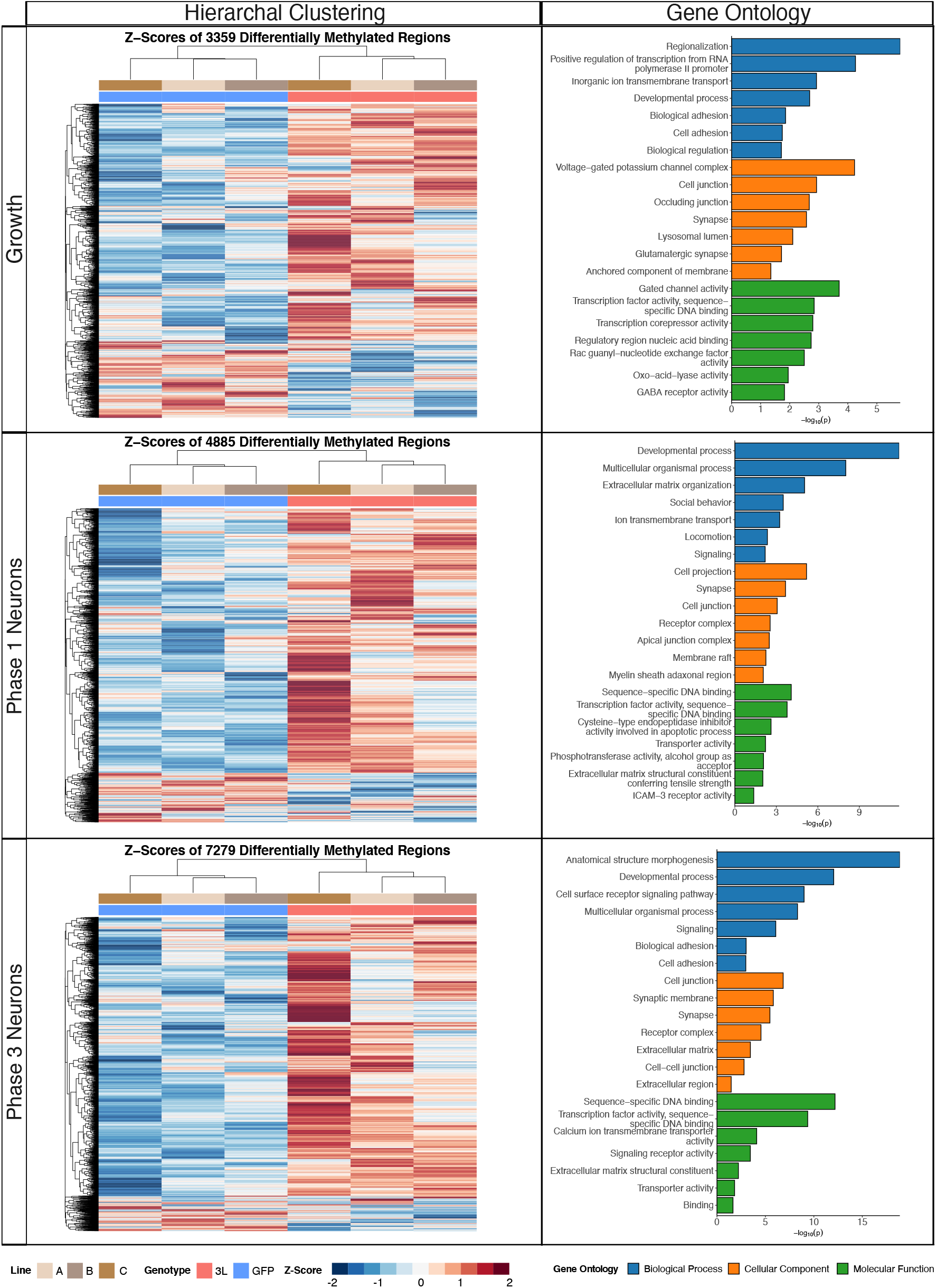
Differentially methylated region (DMR) heatmaps and slimmed gene ontology (GO) enrichments for *DNMT3L* overexpression at the different phases of neural differentiation for all 3 cell line replicates.

**Supplementary Figure 4:**
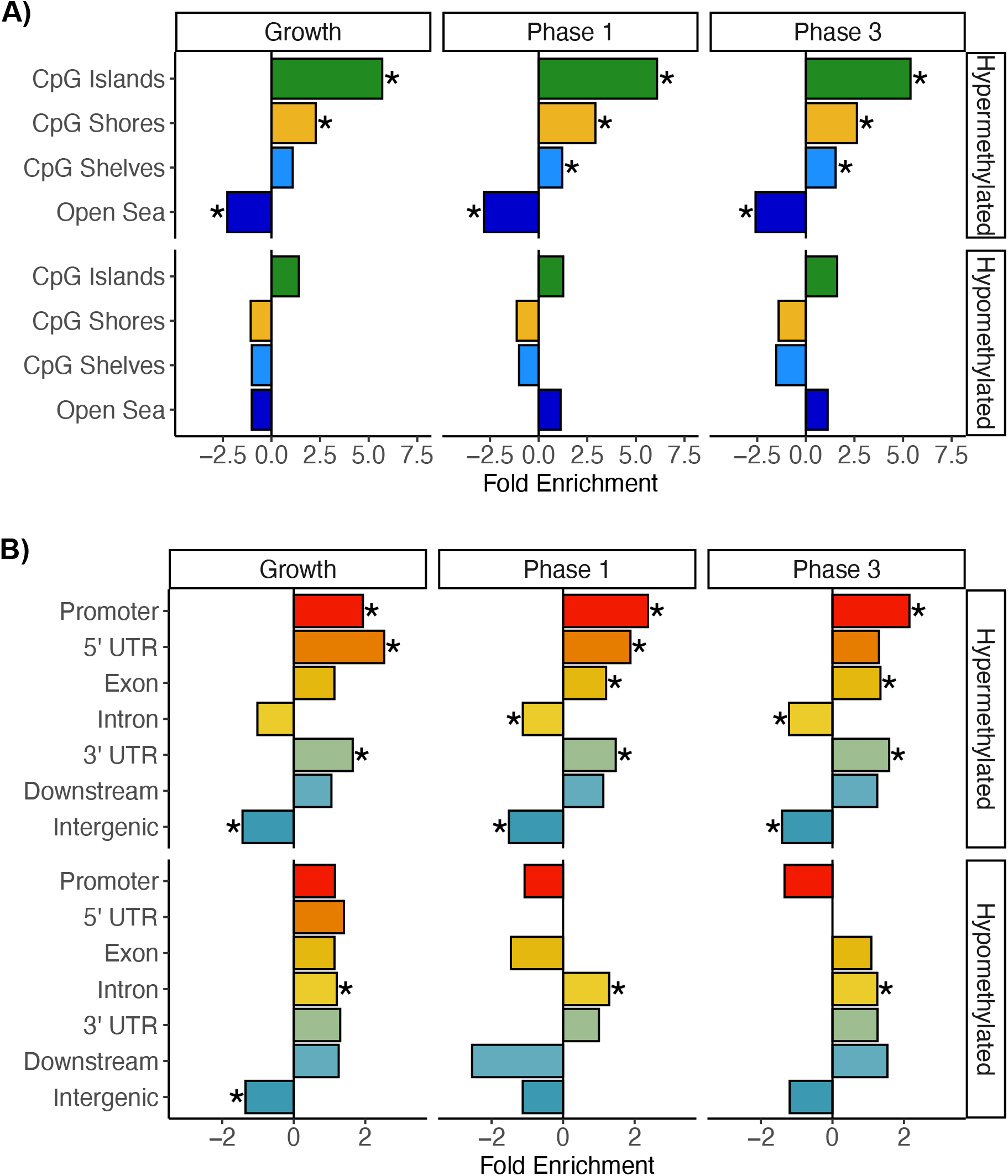
Annotation enrichments for DMRs from the growth, phase 1, and phase 3 comparisons for all 3 cell line replicates. **A)** CpG and **B)** genic annotation enrichments for hypermethylated and hypomethylated DMRs. * = *q* < 0.05.

**Supplementary Figure 5:**
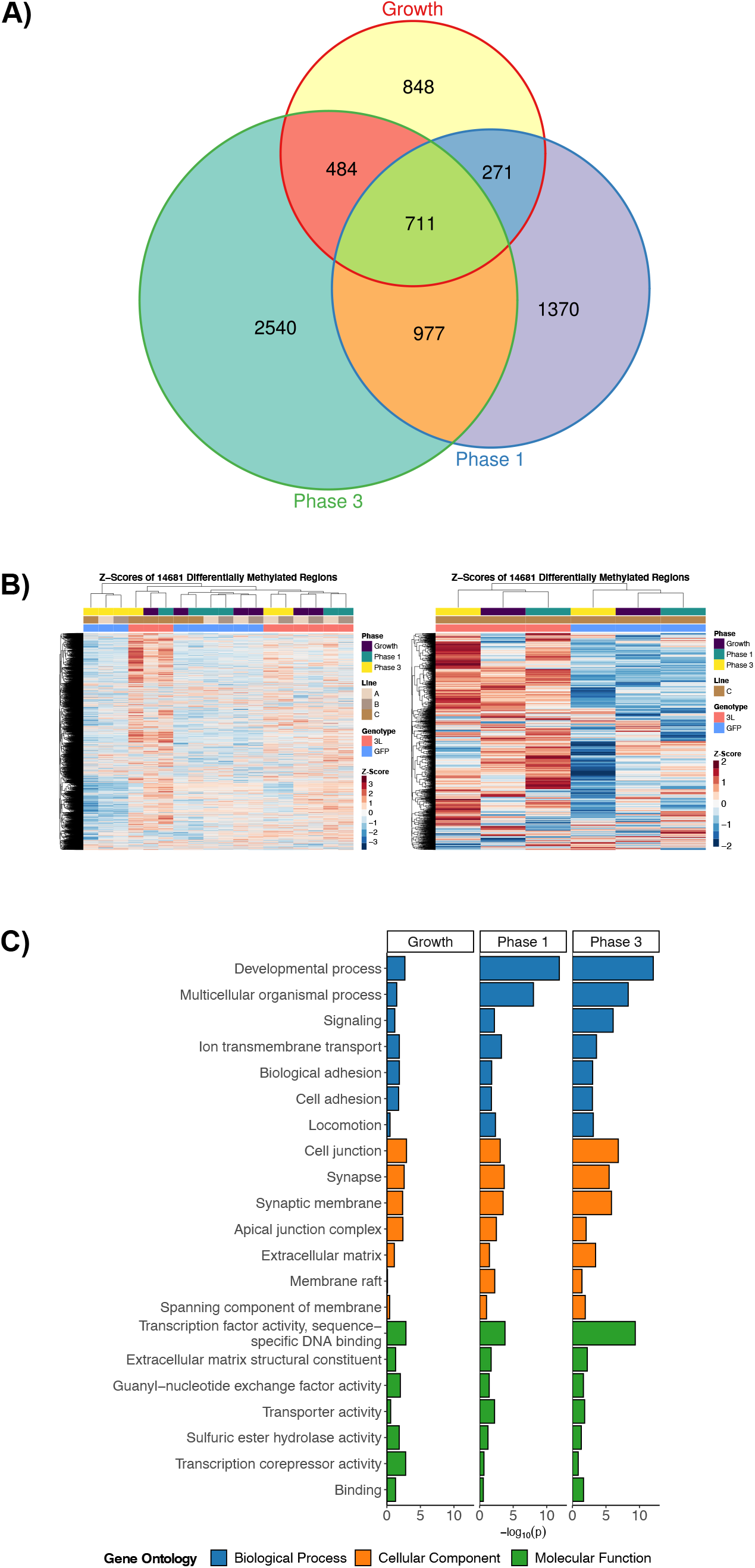
Consensus DMR profiles for *DNMT3L* overexpression across all phases of neural differentiation for all 3 cell line replicates. **A)** Euler diagram of gene symbol overlaps for the genotype comparison at each phase of neural differentiation. **B)** Heatmap of hierarchal clustering of Z-scores for the consensus DMRs that are derived from merging the DMRs from each phase comparison by sequence overlap. **C)** Bar plot of comparison specific p-values from the meta-analysis of the least dispensable significant (*p* < 0.05) slimmed GO enrichments.

**Supplementary Figure 6:**
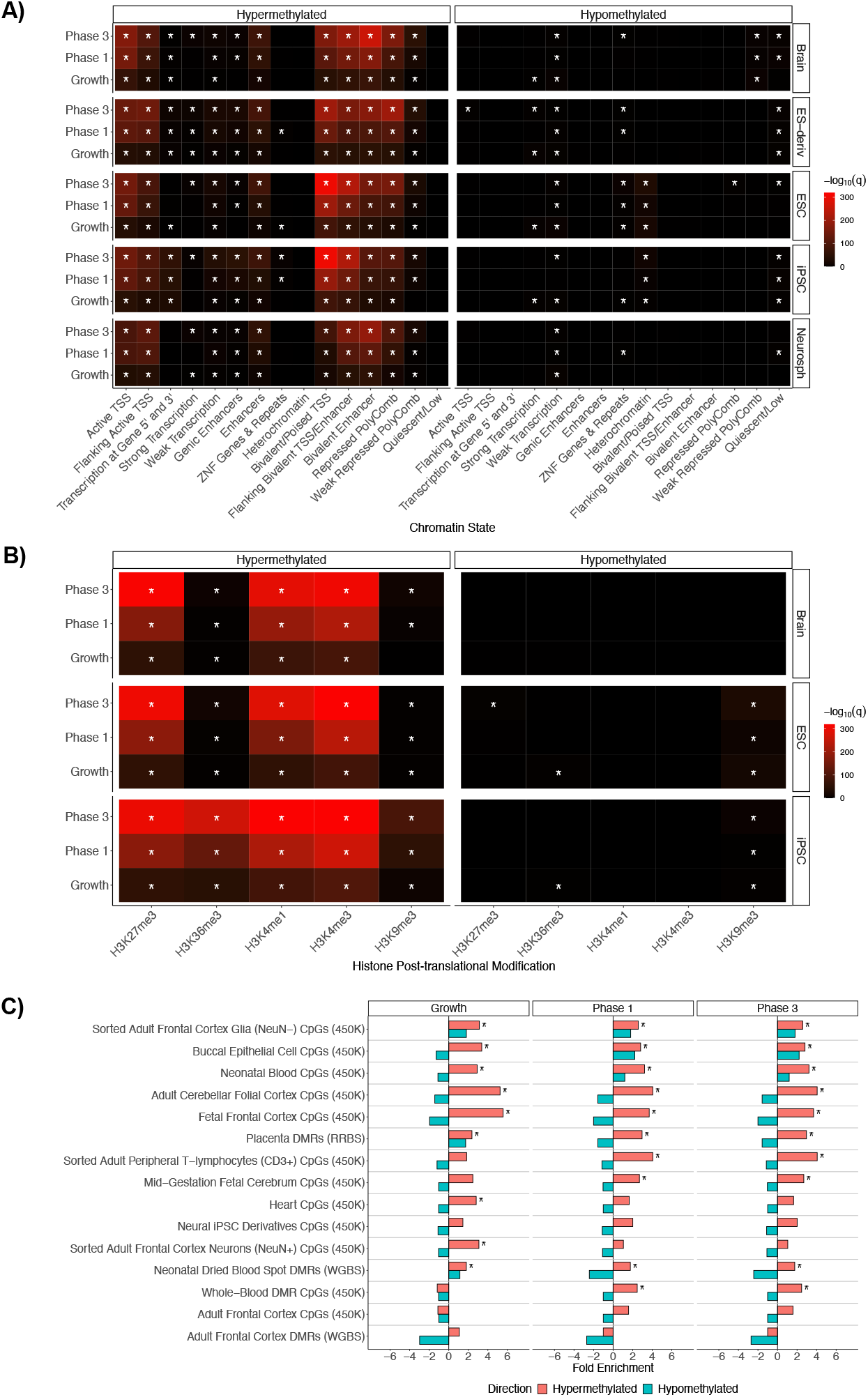
Reference epigenome and cross-tissue Down syndrome enrichment analyses for the hypermethylated and hypomethylated DMRs from the genotype comparisons at each phase of neural differentiation for all 3 cell line replicates. **A)** Summary heatmap of top *q*-values for the chromHMM core 15-state enrichment analyses for the brain, embryonic stem cell derivatives (ES-derivatives), embryonic stem cells (ESC), induced pluripotent stem cells (iPSC), and neurospheres (Neurosph) categories. Values of infinity have been replaced with the top value. **B)** Summary heatmap of *q*-values for Roadmap epigenomics 127 reference epigenomes 5 core histone modification enrichment analyses for the Brain, ESC, and iPSC categories. Values of infinity have been replaced with the top value. **C)** Bar plot of Down syndrome cross-tissue analysis. All enrichments are relative to background regions. * = *q* < 0.05.

**Supplementary Table 1**. Testable background regions and significant DMRs for *DNMT3L* overexpression in **A)** growth phase, **B)** phase 1, and **C)** phase 3.

**Supplementary Table 2**. Testable background regions and significant DMRs for *DNMT3L* overexpression in all 3 cell line replicates for **A)** growth phase, **B)** phase 1, and **C)** phase 3.

